# Male reproductive phenotype and coercive mating performance in the guppy *Poecilia reticulata*

**DOI:** 10.1101/2024.03.10.584275

**Authors:** Alexandra Glavaschi, Elisa Morbiato, Andrea Pilastro

## Abstract

In species with fixed alternative male mating tactics, differences between male phenotypes associated with each tactic are well understood. By contrast, in species with fully interchangeable male mating strategies, associations between male phenotypes and fitness when adopting different tactics have received much less attention. One such species is the Trinidad guppy *Poecilia reticulata,* where males perform high rates of coercive mating attempts (gonopodial thrusts, GTs hereafter) but also switch between GTs and courtship with great flexibility. Male phenotypes favored by females have been described in detail and consist of complex, nonlinear combinations of traits. Coercive tactics also contribute towards male fitness, but no study to date has provided a multivariate description of guppy phenotypes able to obtain fertilizations via GTs, despite evidence suggesting they should be different from phenotypes successful in cooperative mating scenarios. Here we observe male mating behavior in freely interacting mixed-sex groups and compute a GT performance variable based on the closest distance the male approaches the female before abandoning the thrust. We use multivariate selection techniques to relate GT performance to traits and combinations of traits known to contribute towards male fitness. Guppy males that perform best in GTs are small, bold, with large areas of iridescent coloration and fast-swimming sperm, as well as intermediate orange coloration and sperm count. This phenotype only partly confirms our expectation, as it comprises traits advantageous in cooperative mating scenarios. Our study highlights the importance of using multivariate approaches when investigating sexual selection in the context of coercive mating strategies.

## Introduction

Forced copulations are a form of sexual coercion employed by males of a wide range of internally fertilizing species to overcome female unwillingness to mate (Arnqvist and Rowe 2005; Clutton-Brock and Parker 1995). Males rely on forced copulations to various degrees to achieve fertilizations, so that in taxa such as bedbugs *Cimex lectularius* or mosquitofish *Gambusia holbrooki* they constitute the sole mating tactic (Bisazza et al. 2001; Reinhardt and Siva-Jothy 2007) while in other groups like squid *Doryteuthis pleii* and some swordtails *Xiphophorus sp* (Apostólico and Marian 2019; Ryan and Causey 1989) they represent alternative mating strategies, in which some males always rely on sexual coercion, whereas others court females and mate cooperatively (Oliveira et al. 2008). In these cases, males typically differ in their reproductive phenotype: courting/territorial males are usually larger and possess more developed sexual secondary traits, whereas males adopting alternative mating strategies (e.g. coercive, satellite or sneaker males) are often smaller, less conspicuous and invest more in postmating traits, such as sperm number and velocity, due to the high level of sperm competition usually associated with these mating tactics (Lank et al. 2013). Finally, in other species the same individual male can adopt one or the other mating tactic opportunistically, as is the case in damselflies (Cordero and Andrés 2002), or many poeciliid fishes (Evans et al. 2011a). While relationships between male reproductive phenotypes and fixed alternative mating tactics are relatively well understood (Gross 1996; Shuster 2010), the association between a male’s phenotype and his performance in coercive matings in species with fully interchangeable tactics has received markedly less attention. The guppy, *Poecilia reticulata*, is a well-suited organism for investigating the association between a male’s reproductive traits and his performance when attempting coercive matings. Guppies are small freshwater fish native to Central America, with internal fertilization and a polygynandrous mating system and males under strong pre and post-copulatory sexual selection (Devigili et al. 2015). The anal fin in males is modified into an intromittent organ, the gonopodium, used to transfer sperm to females. Males readily alternate between two mating tactics: i) courtship, or sigmoid displays (SDs hereafter), whereby they arch their bodies in an S-shape and display their coloration to females, and ii) sneak mating attempts, or gonopodial thrusts (GTs hereafter), whereby they swing their gonopodium forward and approach the female from behind in an attempt to forcibly inseminate her (Houde 1997; Magurran 2005). Female guppies are generally only receptive as virgins and for 3-4 days following parturition (Liley 1966), when they are likely to respond to male courtship by mating cooperatively. If courtship fails to obtain a cooperative mating, males commonly switch to sneaking (Houde 1997). Given that females are highly selective of mates and have a short receptive window, males devote an extreme amount of time to sneaky mating attempts, perhaps the highest observed for any organism in the wild, with females being subject to up to one GT per minute (Magurran and Seghers 1994b). Only a small fraction of sneaky attempts (possibly less than 2%) result in sperm transfer (Pilastro and Bisazza 1999), with the amount of sperm usually significantly lower than that transferred during cooperative matings (Pilastro and Bisazza 1999; Pilastro et al. 2007). Even if individual sneaky mating attempts are unlikely to lead to successful fertilizations (Russell et al. 2006), this tactic can still contribute to male fitness given the astronomic number of coercive matings attempted throughout a male’s lifespan (Matthews and Magurran 2000) and the fact that female-stored sperm can fertilize eggs several months after copulation (López-Sepulcre et al. 2013; Winge 1937).

Both cooperative and coercive matings are therefore likely to contribute to male guppy reproductive success, but optimal male phenotype at courting is likely to differ from that at sneaking for the following reasons. First, in other poeciliid species where the two mating strategies have a genetic basis, males adopting the coercive tactic are smaller and less conspicuous than courting males (Ryan and Causey 1989; Zimmerer and Kallman 1989). Second, in Trinidadian guppies, males inhabiting high predation sites are smaller, have duller coloration and rely more on sneaky matings than males from low predation localities, which are larger, more conspicuous and court females more frequently (Reznick and Travis 2019). Third, negative associations between body size and ornamentation respectively and GT frequency have been reported in lab experiments: despite the high plasticity of guppy male mating behavior, small drab males perform consistently more GTs compared to their large colorful counterparts, irrespective of the level of predation pressure experienced by the population of origin (Bisazza and Pilastro 1997; Endler 1995; Reynolds et al. 1993).

Guppy male phenotypes that increase attractiveness and reproductive success in cooperative mating scenarios have been studied in detail. While larger and more colorful males with higher courtship rate are usually preferred by females (Houde 1997), several studies have evidenced that multiple pre- and postcopulatory traits, including body size, gonopodium length, areas of coloration, and courtship rate contribute towards male reproductive success through complex, nonlinear interactions (Blows et al. 2003; Brooks and Endler 2001; Cattelan et al. 2020; Devigili et al. 2015; Glavaschi et al. 2022). Sperm reserves and sperm velocity also affect male reproductive success directly in sperm competition (Boschetto et al. 2011) and indirectly, through phenotypic (Locatello et al. 2006) and genetic correlations with precopulatory traits (Evans 2010). Furthermore, high risk-taking propensity is favored by females (Godin and Dugatkin 1996), and associated with a higher reproductive success (Herdegen-Radwan 2019). But whether or not the optimal male phenotype at sneaking is also characterized by similarly complex combinations of traits has never been investigated in guppies. In the closely related eastern mosquitofish *Gambusia holbrooki*, in which males rely exclusively on coercive matings for obtaining fertilizations, multivariate selection analysis revealed predominantly non-directional and correlational associations between male sexual traits and insemination success, highlighting the importance of investigating sexual selection using a multivariate framework in the context of coercive matings (Head et al. 2015).

Here we aim to identify traits (and combinations of traits) predicting forced copulation success of male guppies with sexually unreceptive females by applying multivariate selection analysis followed by canonical rotation. The success rate of coercive mating attempts is generally challenging to estimate accurately: in a typical mating trial, lasting for a maximum of one hour during which male and female behavior can be continuously monitored, the probability of observing a successful sneaky mating is usually exceedingly low, therefore most estimations of coercive mating success would be zero inflated. We therefore used a more continuous measure of male capability to achieve sperm transfer through coercive copulations, originally developed for the eastern mosquitofish (Bisazza and Pilastro 1997). Our estimate of male performance in GT is based on how closely the gonopodium tip approaches the female gonopore in each mating attempt, according to an *a priori* developed scale (see Materials and methods). We expect the optimal phenotype at sneaking to be composed of complex associations between pre and postmating traits, contributing both directly and indirectly towards GT performance.

## Materials and methods

### Fish maintenance

Guppies used in this experiment originate from a stock captured from Lower Tacarigua River (high predation regime) in Trinidad in 2002. Fish are kept in 150L tanks at 25-27°C, at an approximately equal sex ratio, 1 fish/L density and fed dry food (DuplarinS) daily and *Artemia salina* nauplii three times a week. New-born are collected monthly and separated into single-sex tanks as soon as sexual traits become visible, to obtain virgin females and males of known age.

### Experiment overview

A total of ninety-six adult males (aged between four and six months), split into twenty-four replicates of four fish each, were used for this experiment. Data collection timeline is shown in Figure S1A. Fish were selected from stock populations ensuring that within each replicate males were individually recognizable by the human observer according to color patterns. To allow males to replenish sperm reserves while maintaining sexual motivation (Bozynski and Liley 2003), each replicate was isolated for three days in a 38 x 25 x 40 cm (H x L x D) holding tank filled with 20 cm of water, containing gravel, an aerator and two females inside a clear perforated cylinder (10 cm diameter) in the center. Males were kept in holding tanks for the whole duration of data collection except for behavioral observations and boldness tests (see below) for a maximum of eight days for each replicate. We used nonreceptive stimulus females for behavioral observations in order to maximize the likelihood that males would perform GTs, as in freely interacting mixed-sex groups where they can perceive olfactory cues, males direct more coercive mating attempts to this category compared to virgin or post-partum females (Guevara-Fiore et al. 2009). We selected adult females (∼ 20 mm standard length) from mixed-sex stock populations and confirmed pregnancies (and therefore non-receptiveness) by visual examination. Females were allowed to habituate to the testing room for at least three days. Acclimation tanks were checked twice daily and no broods were detected for the duration of the experiment.

Males were subject to two boldness tests, first the day following mating trials and the second two or three days later. After the second boldness test, males were photographed and stripped of sperm for ejaculate analysis. All fish were released into post-experimental tanks following data collection and not reused in other studies.

### Estimates of GT performance

Prior to data collection, we constructed a scale of GT intensity consisting of five levels based on how closely, in theory, could the male approach the female during a sneaky mating attempt. Thus, a level 1 GT indicates a coercive mating attempt abandoned more than a body length away from the female, while a level 5 GT indicates a successful thrust (with sperm transfer) inferred from male postcopulatory jerks (detailed in Table 1).

**Table 1.**
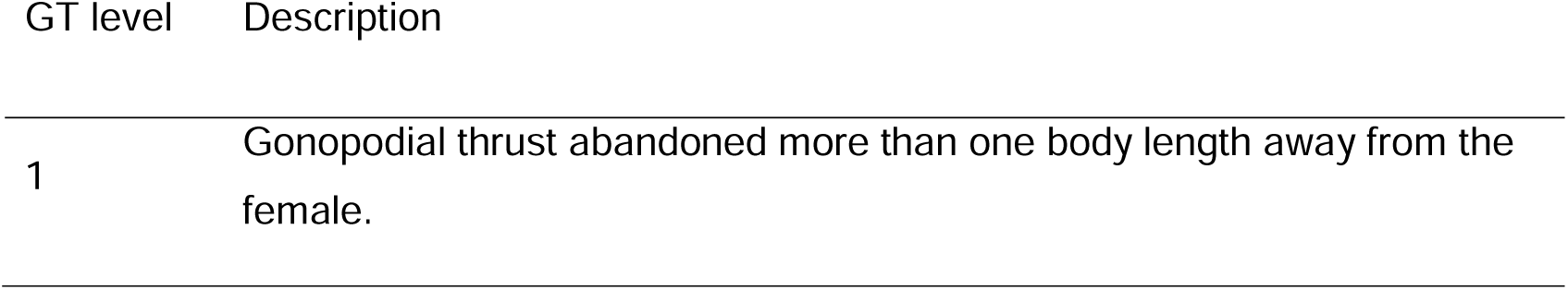
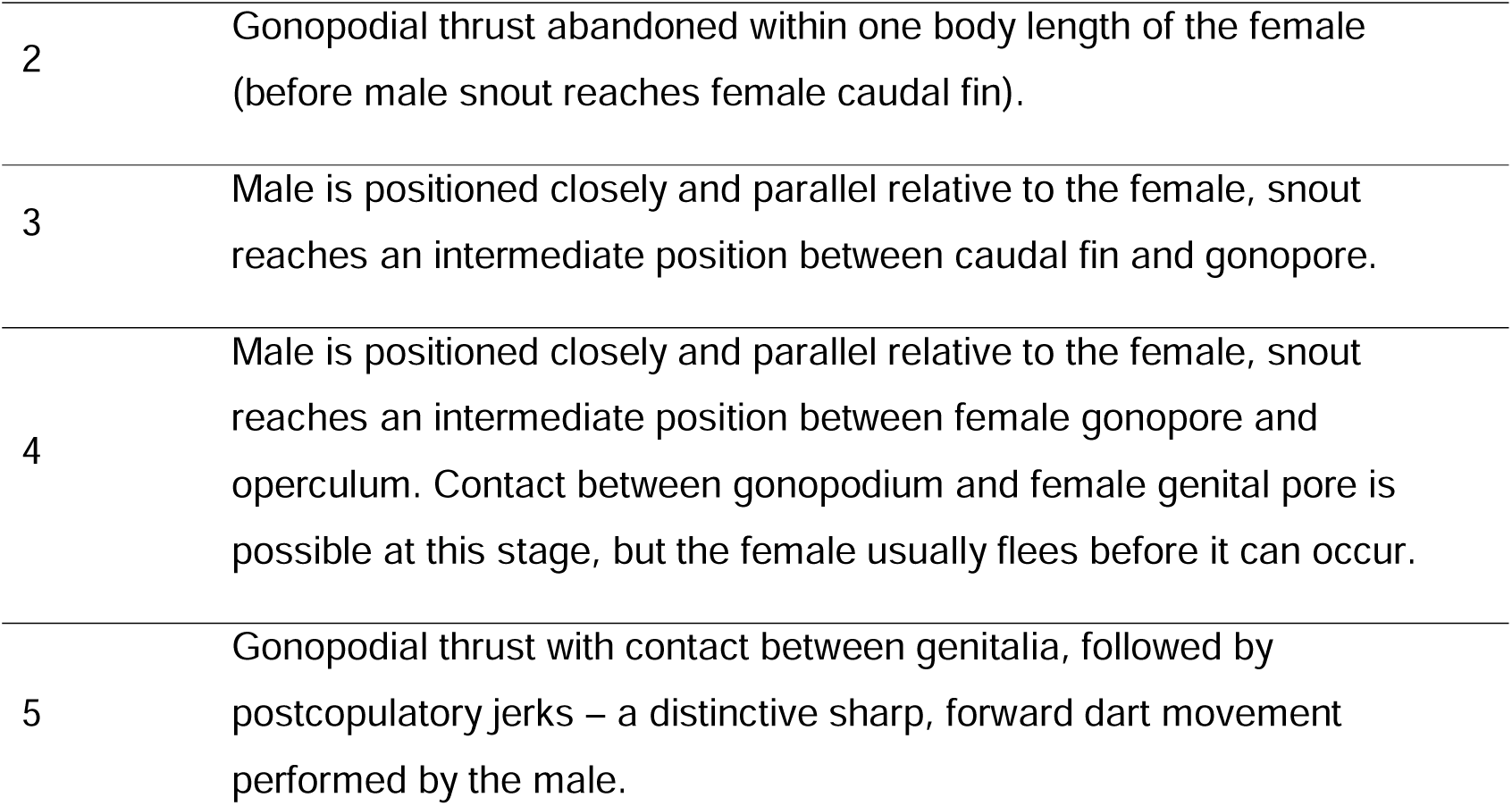
Levels of GT intensity and descriptions. A subset of the attempts scored as 4 could have resulted in a contact between gonopodium and gonopore. However, since this is difficult to confirm in the absence of slow-motion video footage, we conservatively considered a successful forced copulation only that followed by jerking (stage 5, Liley 1966).

Behavioral observations were conducted in 38 x 83 x 40 cm (H x L x D) tanks filled with 20 cm of water containing gravel and an aerator. Opaque removable partitions divided the tanks transversally into two sections, 25 and 58 cm long, respectively (Figure S1B). Twenty-four hours before the trial, six non-receptive females were transferred into the larger section and a male replicate into the smaller section of the tanks. We observed two male replicates per day, between 08:00 and 11:00, when guppies are most active (Magurran 2005). The partition was removed and fish allowed to recover from the disturbance (until normal swimming resumed, usually within two minutes), after which the observation started. One observer conducted sequential focal sampling, following each male for 15 minutes, and recorded the number of GTs and the level of each GT (Table 1), as well as the frequency of SDs. No cooperative matings occurred in any replicates. There was a single instance of GT followed by jerking, and hence presumably with sperm transfer (Figure 1A). We therefore pooled the number of GT attempts that reached level 4 and 5 (Table 1) into a unique index, hereafter referred to as “GT performance”, that we include as dependent variable in the analysis.

**Figure 1.**
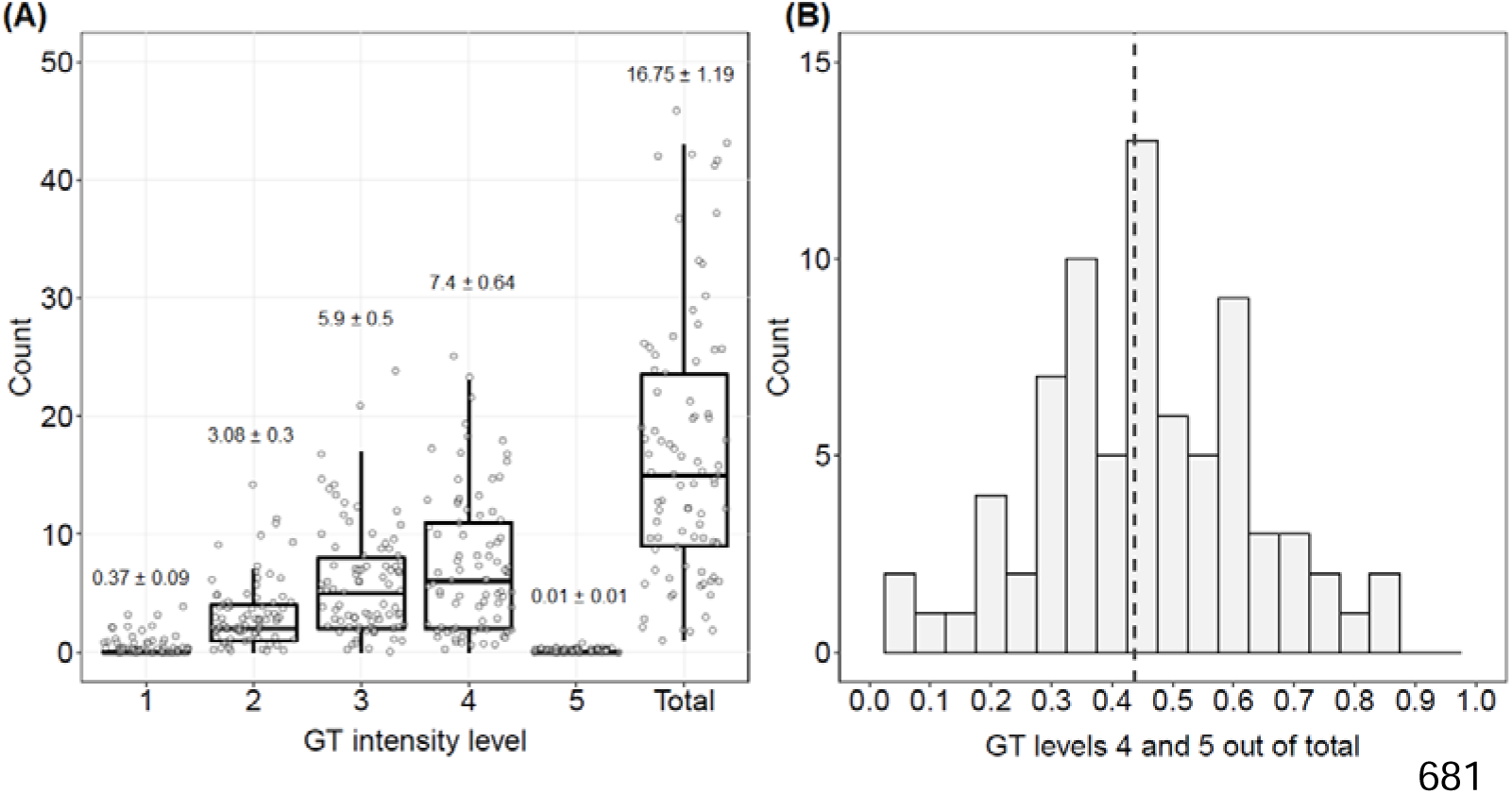
(A) Distribution of GT frequencies per 15 minutes according to each level. Numbers above boxplots represent means ± standard errors. See Table 1 for descriptions of intensity levels. (B) Distribution of the proportion of GTs most likely to result in sperm transfer (levels 4 and 5 pooled, i.e. our measure of GT performance, see text) out of the total. Dashed line represents the sample average of 0.4.

### Trait measurement

Boldness tests were carried out in a white circular arena of 40 cm diameter containing water up to a depth of 2.5 cm, illuminated by two 15-W neon lights on each side. The arena had a refuge in the center, consisting of a plastic circle mounted on three equidistantly placed supports, 4 cm in diameter and 1.5 cm high. A Panasonic HCV180 video camera was mounted 1m above the arena. The fish was released close to the refuge, 200 µl alarm substance (prepared according to (Nordell 1998)) pipetted into the water and its behavior recorded for 10 minutes. Videos were scored using BORIS 7.1.3 ((Friard and Gamba 2016), http://www.boris.unito.it/pages/download.html) and the following behaviors measured: latency to leave the refuge (inverted to facilitate interpretation), total time spent underneath the refuge and number of visits to refuge. We scored as “visit to refuge” any instance of fish returning to the refuge from at least three body lengths away, whether to hide or swim straight through, therefore this measure indicates fish willingness to explore the arena. Each male underwent two boldness tests, separated by 48-72h.

Following the second boldness test, males were anesthetized in an MS-222 bath, placed on a grid-lined slide under a dissection microscope equipped with a Canon 450D camera and their left sides photographed. Fish were stripped of ejaculates by swinging the gonopodium back and forth and applying gentle pressure to the abdomen. In this species, sperm is organized in discrete units (spermatozeugmata or bundles), each containing around 21000 individual cells (Boschetto et al., 2011). Sperm bundles were released into a drop of saline solution and the ejaculate photographed for the purposes of sperm counting. Photographs of males and ejaculates were scored using ImageJ software and the following variables recorded: number of sperm bundles, body area, standard length, tail area, gonopodium length, and areas of orange and iridescent coloration, respectively.

Using a Drummond micropipette, three sperm bundles were transferred to a multi-well slide coated with 1% polyvinyl alcohol to prevent cells from sticking to the glass (Boschetto et al., 2011). Sperm were activated in 3 µl of medium containing 150 mM KCl and 2 mg/L bovine serum albumin (Billard et al.,1990). Velocity was measured using a CEROS sperm tracker (Hamilton-Thorne Research, Beverly, MA, USA) as cells were swimming away from the opening bundle. Of the three measures of velocity provided by the tracker (average path, VAP; curvilinear, VCL; straight line, VSL), we retained VAP for further analyses (Cattelan et al. 2020; Devigili et al. 2019). Sperm velocity for each male was measured from 394 ± 16.3 (mean ± SE) cells.

### Data analysis

After excluding nine males for which we could not obtain measures for all traits of interest and six males that did not perform any sexual behaviors, our final sample size consisted of 81 males. Ejaculate photographs of two males (different from those above) were lost due to equipment failure and for these fish we replaced sperm count values with sample average.

The three measures we collected during the two boldness tests were significantly repeatable within individual (all R > 0.3 according to the formula proposed by (Lessells and Boag 1987)) and were therefore averaged. We conducted a PCA to condense these measures into a single index of boldness. The resulting PC explained 0.69 of the variance and was loaded positively by latency and visits to refuge (factor loadings = 0.93 and 0.72 respectively) and negatively by time spent under refuge (factor loading = -0.83). High PC scores corresponded to fish that had a low latency to leave the refuge, spent less time under the refuge and made more visits to the refuge – an indication of willingness to explore the arena (see above) – and were therefore used as a measure of individual boldness.

To explore the relationship between GT performance and male traits, we ran a linear regression followed by a full quadratic multiple regression, with GT performance (see above and Table 1) as the dependent variable and linear traits, quadratic-transformed traits and all pairwise trait combinations as predictors. The traits considered were (*i*) body area, (*ii*) gonopodium length, (*iii*) area of orange coloration, (*iv*) area of iridescent coloration, (*v*) sperm velocity, (*vi*) sperm number and (*vii*) boldness. Average trait values for each trait are shown in supplementary Table S1. The dependent variable was standardized to a mean of one and predictor variables to a mean of zero and a standard deviation of one (Lande and Arnold 1983). From the linear regression, we obtained linear coefficients (β) equivalent to the slope of the performance surface. We obtained quadratic coefficients (γ) representing the curvature of the performance surface from the full quadratic regression. We doubled quadratic coefficients to account for statistical packages underestimating them by 0.5 (Stinchcombe et al. 2008).

We then conducted a canonical rotation by multiplying the gamma matrix (containing doubled quadratic coefficients on the diagonal and correlational coefficients on the off-diagonals) with the matrix of original trait values (Phillips and Arnold 1989). The new axes of selection so obtained (eigenvectors M1 – M7, Table 3) are characterized by original trait loadings, similarly to a PCA (Blows and Brooks 2003). Each eigenvector obtained by canonical rotation has an associated eigenvalue (λ coefficients). Absolute values of λ coefficients indicate the strength of selection (i.e. curvature of the surface, see below), while the shape of selection is given by their signs: positive λ values correspond to disruptive selection while negative λ values indicate stabilizing selection.

To estimate linear selection (u gradients) along the new axes, we also rotated the original linear coefficients (β) onto the canonical space (Lymbery et al. 2018). We tested the significance of each eigenvector with the permutation procedure suggested by (Lewis et al. 2011). The performance surface built by the new vectors of selection was visualized with the ‘Tps’ function of the ‘fields’ package in R (Nychka et al. 2017). Our analysis can be reproduced using the dataset and code provided as supplementary materials.

### Ethical note

Our data collection protocol was approved by the University of Padova Institutional Ethical Committee (permit no. 256/2018). Sperm stripping and photography was performed by an experienced operator and the entire procedure lasted for less than one minute per male. All fish recovered well from anesthesia and resumed normal swimming within two minutes.

## Results

Figure 1A illustrates the frequency of each level of GT intensity described in Table 1, as well as the total number of GTs across the 81 males observed. The sum of levels 4 and 5, our measure of GT performance, accounted on average for 0.43 of the total GTs attempted (Figure 1B). We did not identify any linear relationships between GT performance and male traits. A positive correlational gradient between iridescence and boldness suggests that males with better GT performance were either bold with a large area of iridescent coloration or shy with a small area of iridescence (Table 2).

**Table 2.** Linear (beta) and nonlinear (gamma) coefficients of the gamma matrix obtained from the second-order multiple regression. Quadratic coefficients are shown on diagonals and indicate disruptive (+) or stabilising (–) relationships with GT performance. Correlational coefficients are shown below diagonals and represent pairs of positively or negatively correlated traits. Significant coefficients (*p* < 0.05) are indicated in bold.

| Gamma coefficients |  |  |  |  |  |  |  |
| --- | --- | --- | --- | --- | --- | --- | --- |
|  | Body area | Gonopodium length | Orange | Iridescent | Sperm velocity | Sperm number | Boldness |
| Beta coefficients | 0.108 | -0.156 | 0.046 | -0.018 | 0.047 | 0.070 | 0.050 |
| Body area | 0.064 |  |  |  |  |  |  |
| Gonopodium length | -0.131 | 0.098 |  |  |  |  |  |
| Orange | -0.062 | 0.140 | -0.207 |  |  |  |  |
| Iridescent | -0.190 | 0.244 | 0.044 | 0.222 |  |  |  |
| Sperm velocity | -0.135 | -0.009 | -0.078 | 0.327 | -0.056 |  |  |
| Sperm number | -0.024 | -0.028 | 0.055 | 0.071 | 0.044 | -0.301 |  |
| Boldness | -0.095 | 0.004 | -0.018 | **0.417** | 0.050 | 0.038 | 0.103 |

Two of the seven axes of selection revealed by canonical rotation had significant λ coefficients. Axis M1 described disruptive selection (positive λ) and was loaded by body area (positive), iridescence (negative), sperm velocity (negative) and boldness (negative). Axis M6 was associated with stabilizing selection (negative λ) and was primarily loaded by sperm number (negative), followed by orange (positive) (Table 3). The fitness surface built by these two axes displays an asymmetric saddle shape (Figure 2). The area of highest GT performance corresponded to the peak at the negative extreme of axis M1 (Figure 2). Thus, guppy males with the highest GT performance scores were small, bold, with large areas of iridescent coloration and fast-swimming sperm, as well as intermediate orange coloration and sperm count. Disruptive selection along the M1 axis suggested the presence of an alternative phenotype at the positive end, consisting of large, shy males, with small areas of iridescence, slow-swimming sperm and intermediate areas of orange and sperm count (Table 3, Figure 2). Note, however, that the fitness surface was characterized by one well-defined peak only, at the negative extreme of M1 where the highest GT performance was concentrated (Figure 2), thus we will restrict our discussion to the former phenotype.

**Figure 2.**
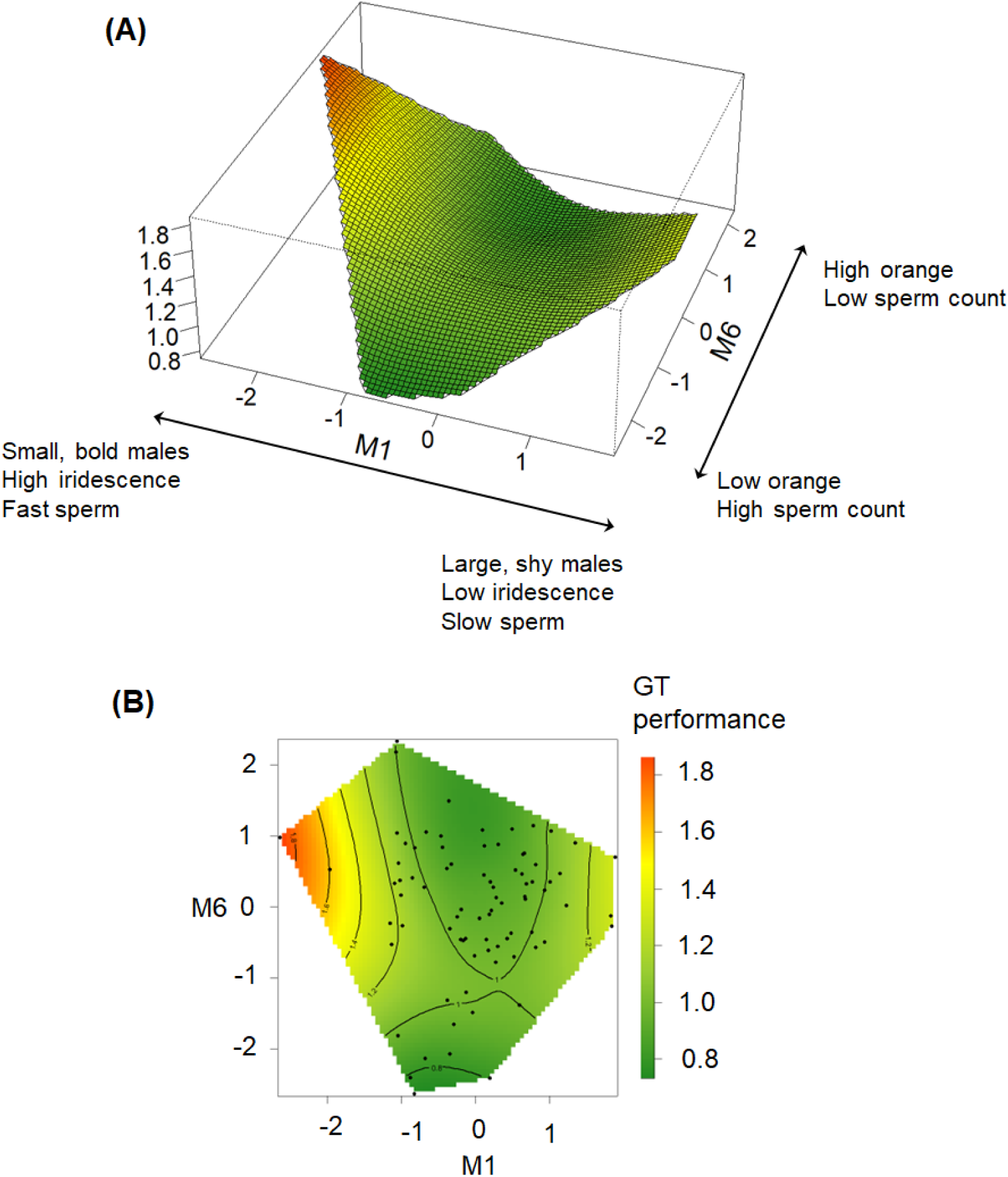
Performance surface (A) and 2D contour plot (B) determined by the two significant axes identified by canonical rotation, M1 (disruptive) and M6 (stabilizing) showing relationships between GT performance and male traits. The peak at the negative extreme of axis M1 corresponds to the phenotype associated with the highest GT performance, consisting of small body size, high boldness, large area of iridescence, fast sperm and intermediate sperm number and area of orange coloration.

**Table 3.** Canonical rotation of the gamma matrix. Eigenvectors obtained by canonical rotation of the gamma matrix and estimates of linear (theta) and nonlinear (lambda) performance gradients. Trait loadings on each eigenvector can be interpreted similarly to those obtained by a principal component analysis. Absolute lambda values reflect the curvature of the surface, while their signs indicate the shape (positive=disruptive; negative=stabilizing). Significant lambda values (*p* < 0.05) are indicated in bold. *P* values obtained with the permutation procedure (5000 iterations) proposed by Lewis et al. 2011.

|  | Theta<br>(*p* value) | Lambda<br>(*p* value) | Body<br>area | Gonopodium<br>length | Orange | Iridescent | Sperm<br>velocity | Sperm<br>number | Boldness |
| --- | --- | --- | --- | --- | --- | --- | --- | --- | --- |
| **M1** | 0.048<br>(0.697) | **0.849**<br>**(0.009)** | 0.334 | -0.290 | -0.060 | -0.696 | -0.321 | -0.074 | -0.456 |
| M2 | 0.163<br>(0.07) | 0.181<br>(0.386) | 0.268 | -0.755 | -0.359 | 0.099 | 0.226 | 0.056 | 0.406 |
| M3 | -0.027<br>(0.752) | 0.01<br>(0.979) | -0.573 | -0.239 | -0.162 | -0.094 | 0.567 | 0.033 | -0.507 |
| M4 | 0.014<br>(0.867) | -0.085<br>(0.612) | 0.677 | 0.284 | -0.175 | 0.269 | 0.455 | -0.044 | -0.387 |
| M5 | -0.127<br>(0.166) | -0.224<br>(0.16) | -0.147 | 0.267 | -0.693 | -0.111 | -0.087 | -0.622 | 0.143 |
| **M6** | 0.018<br>(0.834) | **-0.365**<br>**(0.029)** | 0.045 | -0.261 | 0.543 | 0.127 | 0.129 | -0.776 | -0.031 |
| M7 | -0.037<br>(0.742) | -0.445<br>(0.158) | -0.082 | -0.260 | -0.190 | 0.630 | -0.541 | -0.019 | -0.447 |

## Discussion

Here we examined the relationships between multiple male traits and a proxy of sperm transfer success through coercive matings in guppies. Our experimental design represents a compromise between ecological relevance (freely interacting groups of male and female guppies) and precise estimates of sperm transfer through GTs by individual males. This would require slow-motion video recordings and examining female reproductive tract following staged interactions between a single male and a single female (Magris et al. 2020; Pilastro et al. 2007) Tank dimensions required for the natural expression of behaviors in groups of fish (supplementary Figure S1), in contrast, precluded obtaining high resolution videos. For this reason, data collection relied on live observations of multiple freely interacting fish, which means that our measure of GT “success” (GT levels 4+5, Table 1) realistically contains GTs with contact between male and female genitalia as well as without. In order to successfully transfer sperm during a GT males need to get as close as possible to the female. Sexually unreceptive females, such as those used in this study, usually flee from approaching males and most sneaky mating attempts are therefore aborted at stages 1-3 (see Table 1). Thus, although we could not obtain an estimation of the proportion of GT attempts that achieved sperm transfer, we are confident that our measure of GT performance reflects males’ ability to forcefully inseminate females.

While a single GT has an exceedingly low probability to result in sperm transfer, its low energetic cost compared to courtship displays allows males to adopt this tactic with a very high frequency (Magurran and Seghers 1994a), such that a substantial proportion of nonreceptive females from wild populations presented sperm within their reproductive tracts that could only have originated from coercive matings (Evans et al. 2003; Matthews and Magurran 2000). Given the capability of female-stored sperm to fertilize eggs for extended periods of time (Devigili et al. 2015; López-Sepulcre et al. 2013), the contribution of GTs towards male lifetime reproductive success is likely to be non-negligible. It is therefore important to understand the strength and the shape of sexual selection on male traits when this alternative mating tactic is adopted.

If female choice and male sexual coercion impose different pressures on guppy male phenotypes (Becher and Magurran 2004), we should expect pre-mating traits to be related to our measure of GT performance in opposite ways compared to cooperative mating/reproductive success in this population (Devigili et al. 2015, Cattelan et al. 2020; Glavaschi et al. 2022) and others (Blows et al. 2003; Brooks and Endler 2001; Gordon et al. 2015). More specifically, body size and male color, which are positively associated with mating success when females are sexually receptive, should be negatively associated with coercive mating success with unreceptive females.

We found that males that achieve the highest GT performance are small and bold, have large areas of iridescence, fast sperm and intermediate sperm count and areas of orange coloration (Figure 2), while gonopodium length was not significantly associated with GT performance. Our findings offer only partial support to the idea that successful coercive matings in this species are associated with combinations of traits that are different from those advantageous in a cooperative mating scenario. Although the association between GT performance and these male traits emerged only in the correlational, multivariate analyses, for the sake of clarity we will discuss our results for each male trait singularly.

We found that small males performed better in GTs (Figure 2), in agreement with earlier results from guppies and other poecilids (Bisazza and Pilastro 1997; Pilastro et al. 1997). From this point of view, as expected, sexual selection associated with coercive mating tactics was opposite to sexual selection associated with courtship mating, as large males are usually preferred by females (Houde 1997; Magurran 2005). Small males are less conspicuous and more maneuverable (Bisazza and Pilastro 1997; Pilastro et al. 1997) and therefore may be better able to closely approach the female without being detected. A greater efficiency of small males in achieving sneaky matings may explain why genetically small males are maintained in natural populations of many poeciliids, in spite of the large size advantage in both intrasexual competition and female choice.

We showed that male boldness measured as risk-taking in the novel arena is also positively related to GT performance (Figure 2). Boldness broadly refers to an animal’s propensity to take risks (Réale et al. 2007). Male guppies scoring high on this behavioral axis are advantaged in multiple contexts: they survive longer in the presence of a predator (Smith and Blumstein 2010), are preferred by females in cooperative mating scenarios (Godin and Briggs 1996) and sire more offspring (Herdegen-Radwan 2019). We cannot confirm here if the same facet of risk-taking propensity is both attractive to females and advantageous in sneaky mating attempts, or if different components of an underlying “boldness construct” are advantageous in different contexts. For example, females are larger than males and can respond aggressively to sneaky mating attempts (Bruce and White 1995; Gorlick 1976), and bold males may be less intimidated by female reaction. Alternatively, if bold males are in better condition, as shown for other poeciliids (Brown et al. 2007), hence faster swimmers, they could be advantaged in a sneaky mating attempt, in which case the relationship between GT performance and boldness is an indirect one.

We also revealed unexpected relationships between male traits and GT performance. First, we found that colorful males, with large areas of iridescence and intermediate areas of orange, achieved the highest levels GT (Figure 2). This combination of traits is also associated with reproductive success during cooperative matings in this same guppy population (Devigili et al. 2015; Glavaschi et al. 2022). Colorful males are expected to be more conspicuous and females should evade their coercive mating attempts more easily, contrary to our result. We can only speculate on the reasons why conspicuous males performed better in GTs. One possibility is that colorful males may swim faster (Nicoletto 1991), a characteristic that should be advantageous when rapidly approaching the female from behind. Alternatively, the iridescent coloration could hinder a female’s ability to pinpoint the male’s exact position, in particular when several males and females interact simultaneously, similarly to what has been suggested to occur in prey-predator systems (Pike 2015).

Second, we found that males with intermediate rather than large sperm reserves display the highest GT performance (Table 2; Figure 2). If males adjust reproductive effort according to current sperm reserves (Matthews et al. 1997), then we can expect larger sperm counts to be positively associated with GT performance. Interestingly, a similar stabilizing selection on the size of sperm reserves has also been found in studies based on cooperative matings in guppies (Cattelan et al. 2018; Glavaschi et al. 2022) and in mosquitofish, where males rely on forced matings to obtain fertility (Head et al. 2015). Stabilizing selection on sperm number in guppies is determined by sexual selection and is attributable to trade-offs with other traits (Cattelan et al. 2018), while in mosquitofish the apparent disadvantage of males with large sperm reserves can be explained as a by-product of the correlation between sperm number and another unmeasured trait related to insemination success (Head et al. 2015). The “missing trait problem” is indeed common to all selection/performance analyses (Morrissey et al. 2012) including ours, because 1) one can never be certain to have included all traits associated with fitness/performance and 2) they are correlational and any observed relationships between traits and fitness/performance do not necessarily indicate causation. The second point is perhaps best illustrated here by the contribution of sperm velocity to the phenotype associated to high GT performance: males succeed in closely approaching females not because of having fast-swimming sperm, but most likely due to another advantageous trait positively correlated with sperm velocity.

Finally, we found no association between gonopodium length and GT performance. This was surprising, since it makes intuitive sense for longer gonopodia to be advantageous in sneaky mating attempts, an idea supported by earlier work (Evans et al. 2011b). While a long gonopodium is probably advantageous in coercive sperm transfer, our measure of GT performance is likely more closely associated with approaching the female without being detected, rather than with successful insemination. Furthermore, even if males with longer gonopodium may “specialize” in GTs, a longer gonopodium may decrease swimming performance and reduce r efficiency in the initial part of a GT attempt (Langerhans et al. 2005). Thus, the two components of a GT attempt, i.e. approaching the female without being detected, and sperm transfer success may exert opposite selective pressures on gonopodium length and shape (Evans et al. 2011b; Kwan et al. 2013).

Here we identified predictors of GT performance in guppies in standard, benign laboratory conditions, but the adoption of this mating behavior in guppies is sensitive to variation in a wide range of environmental factors including: predation risk (Evans et al. 2002; Glavaschi et al. 2022; Godin 1995), sex ratios (Řežucha and Reichard 2014), population density and sex ratio (Magris et al. 2018), light intensity (Chapman et al. 2009), food deprivation (Cattelan et al. 2020), chemical contaminants (Fursdon et al. 2019) and high temperature (Breedveld et al. 2023). Therefore, it may be interesting to investigate whether male phenotypes associated with GT performance under different environmental conditions will differ from the ones identified in the present experiment, as is the case for phenotypes favored by females (Cattelan et al. 2020; Glavaschi et al. 2022).

To the best of our knowledge, our study on guppies and that of Head et al. 2015 on mosquitofish are the only attempts so far to examine relationships between multiple male traits and their combinations and coercive mating success. Although coercion is the most common male mating behavior observed in this fish family and the exclusive mating tactic in approximately half of the species (Reznick et al. 2021), most attention has been directed towards the role of female choice on the evolution of male reproductive phenotype (Evans et al. 2011a). Our understanding of the strength and shape of sexual selection on male phenotypes associated with gonopodial thrusting is therefore still very limited, in particular for species in which males can adopt both tactics opportunistically. In these species it is logistically difficult to distinguish fertilizations deriving from cooperative matings from those following coercive matings. This is also because, even when males do not court and only rely on coercion, there is evidence that females may indirectly favor certain male phenotypes over others (Bisazza et al. 2001). Our approach certainly has limitations, as discussed above. However, it may still constitute a feasible method to investigate selective pressures on male phenotypes associated with sexual coercion in this fish family.

## Supplementary materials

**Figure S1.**
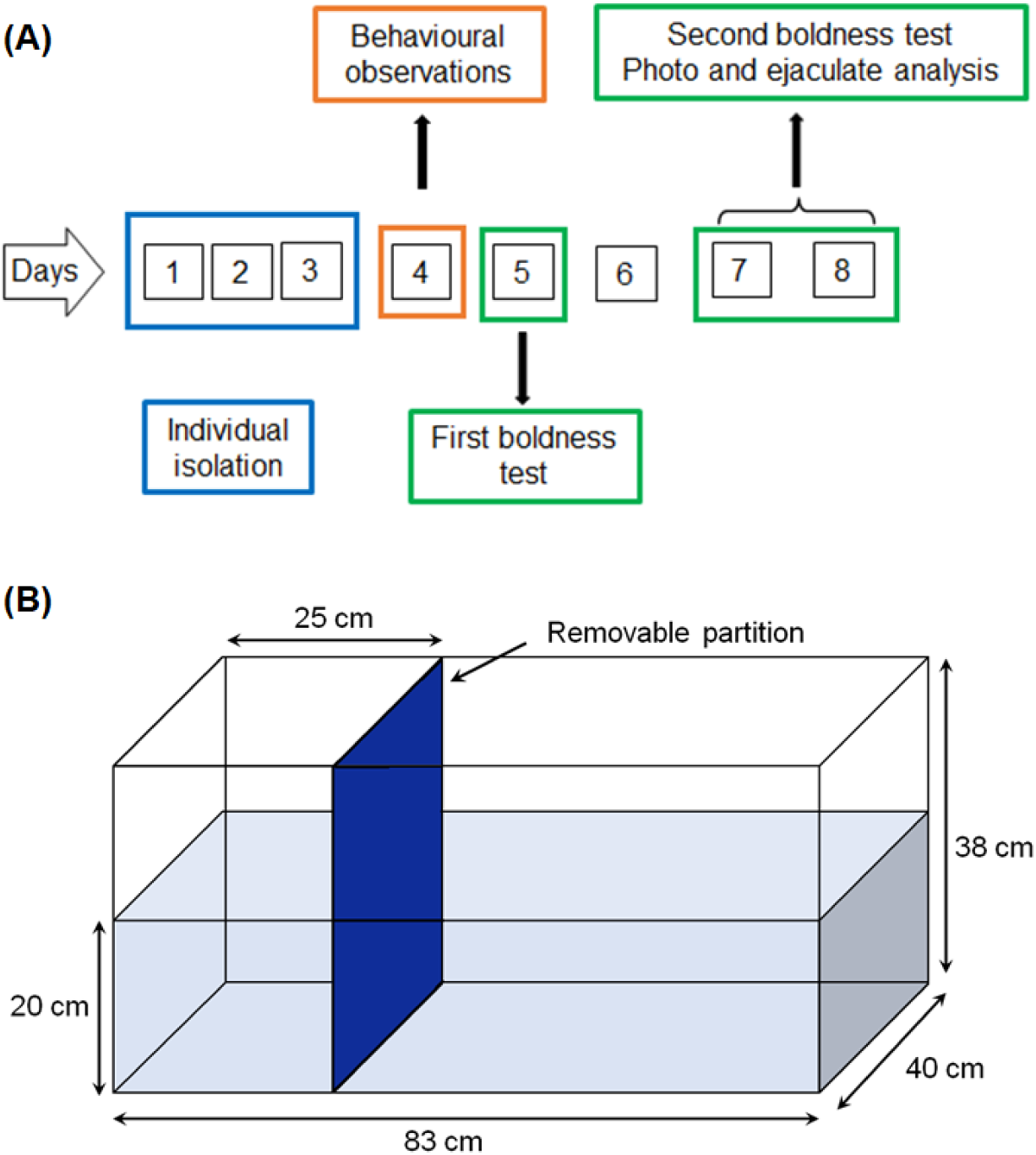
A) Data collection sequence for one male replicate. B) Schematic representation of experimental tank used for behavioral observations (not to scale). Fish were habituated to the tank for 24h, with females in the larger section and males in the smaller.

**Table S1.** Replicate and total averages ± standard deviations of traits used in the analysis. Boldness is a principal component of number of visits to refuge, time spent in refuge and latency to leave refuge.

| Replicate | Body area | Gonopodium | Orange | Iridescent | Sperm velocity | Sperm number | Boldness |
| --- | --- | --- | --- | --- | --- | --- | --- |
| M01 | 44.83 ± 3.59 | 4.25 ± 0.56 | 6.5 ± 3.39 | 3.66 ± 3.37 | 88.02 ± 9.59 | 1.95 ± 0.94 | -0.14 ± 1 |
| M02 | 54.83 ± 4.45 | 3.92 ± 0.32 | 3.6 ± 1.79 | 6.26 ± 1.11 | 93.24 ± 12.45 | 2.81 ± 1.07 | -0.93 ± 0.94 |
| M03 | 51.83 ± 7.36 | 3.91 ± 0.33 | 7.42 ± 3.75 | 6.09 ± 1.67 | 89.32 ± 4.73 | 3.4 ± 2.61 | 0.16 ± 0.63 |
| M04 | 50.17 ± 10.25 | 3.84 ± 0.33 | 6.9 ± 1.37 | 7.64 ± 3.52 | 96.74 ± 20.03 | 3.19 ± 1.26 | 0.15 ± 1.29 |
| M05 | 44.62 ± 9.02 | 3.55 ± 0.16 | 9.73 ± 5.43 | 5.79 ± 3.3 | 82.46 ± 1.44 | 1.49 ± 0.37 | 0.82 ± 1.04 |
| M06 | 46.09 ± 5.55 | 3.76 ± 0.44 | 6.63 ± 7.61 | 10.88 ± 4.16 | 97.43 ± 2.6 | 6.06 ± 3.63 | -0.9 ± 0.82 |
| M07 | 52.82 ± 2.99 | 3.58 ± 0.4 | 8.82 ± 5.49 | 8.73 ± 4.14 | 85.51 ± 12.12 | 4.42 ± 3.74 | 0.54 ± 0.92 |
| M08 | 48.2 ± 8.01 | 3.6 ± 0.26 | 18.43 ± 5.89 | 6.65 ± 4.37 | 93.24 ± 6.41 | 5.23 ± 1.38 | -0.28 ± 1.55 |
| M09 | 50.23 ± 5.03 | 3.81 ± 0.23 | 9.35 ± 5.48 | 7.06 ± 1.06 | 78.37 ± 3.13 | 3.8 ± 3.84 | 0.51 ± 0.64 |
| M10 | 49.01 ± 2.92 | 3.95 ± 0.32 | 14.97 ± 4.39 | 8.29 ± 4.57 | 71.4 ± 5.54 | 3.08 ± 1.56 | -0.85 ± 1.28 |
| M11 | 48.19 ± 4.48 | 3.62 ± 0.35 | 7.58 ± 3.87 | 5.42 ± 2.01 | 78.44 ± 5.41 | 2.11 ± 1.86 | 0.77 ± 0.67 |
| M12 | 46.22 ± 2.48 | 3.71 ± 0.29 | 9.28 ± 3.64 | 7.1 ± 1.35 | 89.84 ± 4.19 | 1.24 ± 0.68 | 0.49 ± 0.47 |
| M13 | 57.26 ± 11.2 | 3.7 ± 0.33 | 19.28 ± 6.56 | 9.85 ± 6.25 | 77.91 ± 8.72 | 6.58 ± 1.12 | 0.81 ± 0.28 |
| M14 | 47.62 ± 4.41 | 3.86 ± 0.19 | 14.58 ± 5.26 | 8.79 ± 1.39 | 82.94 ± 8.07 | 3.32 ± 1.5 | 0.02 ± 0.46 |
| M15 | 48.66 ± 1.92 | 3.6 ± 0.21 | 17.11 ± 13.69 | 8.06 ± 4.2 | 87.26 ± 5.34 | 2.82 ± 0.1 | 0.92 ± 0.86 |
| M16 | 44.11 ± 5.62 | 3.57 ± 0.31 | 6.91 ± 5.11 | 8.22 ± 1.77 | 79.83 ± 6.31 | 5.42 ± 1.95 | -0.14 ± 1 |
| M17 | 44.53 ± 4.66 | 4.03 ± 0.22 | 8.19 ± 2.51 | 6.7 ± 4.11 | 81.23 ± 3.45 | 2.16 ± 1.15 | -0.91 ± 1 |
| M18 | 51.88 ± 2.5 | 4.02 ± 0.14 | 10.13 ± 4.04 | 7.62 ± 3.51 | 79.76 ± 5.87 | 2.67 ± 0.91 | 0.16 ± 0.84 |
| M19 | 45.36 ± 2.09 | 4.03 ± 0.61 | 13.01 ± 7.57 | 5.49 ± 0.17 | 79.21 ± 2.88 | 6.5 ± 4.54 | 0.2 ± 0.94 |
| M20 | 47 ± 5.6 | 3.79 ± 0.33 | 14.33 ± 4.94 | 6.02 ± 4.05 | 80.85 ± 4.38 | 3.9 ± 2.55 | 0.14 ± 0.75 |
| M21 | 43.54 ± 4.33 | 4.51 ± 0.79 | 9.71 ± 2.36 | 5.87 ± 1.77 | 74.06 ± 4.81 | 3.62 ± 2.01 | -0.25 ± 0.54 |
| M22 | 44.55 ± 5.47 | 4.21 ± 0.07 | 11.45 ± 4.45 | 3.21 ± 2.37 | 72.32 ± 2.88 | 2.64 ± 1.85 | -0.02 ± 1.04 |
| M23 | 51.44 ± 4.35 | 3.99 ± 0.51 | 11.84 ± 4.64 | 4.27 ± 1.73 | 90.56 ± 13.02 | 2.12 ± 1.03 | -1.19 ± 1.8 |
| M24 | 47.8 ± 3.44 | 4.2 ± 0.42 | 14.1 ± 3.11 | 5.16 ± 1.67 | 72.62 ± 7.44 | 5.09 ± 2.05 | 0.13 ± 0.75 |
| Total | 48.39 ± 5.85 | 3.88 ± 0.4 | 10.87 ± 5.72 | 6.79 ± 3.18 | 83.29 ± 10.25 | 3.54 ± 2.29 | -0.02 ± 1.01 |

